# Weight locking as a safeguard for open-weight biological foundation models

**DOI:** 10.64898/2026.07.07.736795

**Authors:** Aris Karatzikos, Aggeliki Vasilopoulou, Candace SY Chan, Ioannis Mouratidis, Ilias Georgakopoulos-Soares

**Affiliations:** Division of Pharmacology and Toxicology, College of Pharmacy, The University of Texas at Austin, Dell Paediatric Research Institute, Austin, TX, USA; Department of Computer Science, College of Natural Sciences, The University of Texas at Austin, Austin, TX, USA

## Abstract

Biological foundation models can accelerate research, but their open-weight release raises biosecurity concerns. A common safeguard excludes eukaryotic viral genomes from training, but fine-tuning on public viral data reverses it. We confirm that fine-tuning improves held-out viral language modelling, and that the gain comes from the viral data itself. Weight locking by spectral deformation, proposed to prevent fine-tuning, is untested in biological models. We applied it to two Evo-family models and the protein model ESM3-open, against attackers of increasing knowledge. The lock defends against the naive attacker, whose fine-tuning drives capability below pretrained on all three models. It does not defend against an informed attacker, who freezes or fuses the compensation matrix to restore near-unlocked performance. A compensation-free lock we build from the network’s own rescaling symmetries again defeats the naive attacker, but canonical rebalancing removes it without knowledge of the lock. The barrier is one of knowledge, not compute.

---

Genomic foundation models have reached a level of biological capability that makes their open-weight release a concrete biosafety concern (Sandbrink, 2023; Pannu et al., 2025; Soice et al., 2023). Evo (Nguyen et al., 2024) is a 7-billion-parameter foundation model pretrained on roughly 300 billion nucleotides of prokaryotic and phage sequence, and Evo-2 (Brixi et al., 2026) extends genome-scale pretraining across all domains of life. Although eukaryotic viruses were excluded from the pretraining of both models, these models carry information about virological properties of such viruses, including host tropism, pathogenicity and transmissibility, as measured by the Human Virome Understanding Evaluation (HVUE) benchmark (Dutta et al., 2026). These properties are relevant to pathogen design. The standard biosafety response, filtering pathogen sequences from pretraining data, is insufficient, because fine-tuning Evo-2 on genomes of human-infecting viruses lowers its loss on held-out viral sequences and rescues misuse-relevant predictive capabilities (Black et al., 2025; Wei et al., 2025). We confirm this on both models, and show that the gain comes from the eukaryotic viral data itself. Fine-tuning on a corpus of 544 viral genomes measurably improves an unlocked model’s held-out viral language modelling and its downstream virological discrimination in about 2 GPU-hours on a single data-centre GPU (Supplementary Table 1), an attack available to anyone who can rent one.

The field currently lacks a principled technical mechanism between full weight release and complete withholding. Capability locking via spectral deformation (SpecDef) (Rosati et al., 2025) is a candidate for that middle ground. Before release, selected singular values of target weight matrices are inflated by a factor *α* and paired with a compensation matrix that preserves the forward pass exactly. The released model behaves identically to the original, but the loss landscape is ill-conditioned. Curvature with respect to the compensation matrix grows as *α*^2^, so training it is stable only at learning rates too small to learn much within a realistic budget (Rosati et al., 2026).

The governance question is therefore whether this obstruction is strong enough to defend against an informed attacker. Rosati et al. (2026) prove a fundamental limit of this technique, showing that a white-box attacker can expand locked layers into chains of additional linear factors that reduce per-factor spectral norm at linear cost in model size (their Theorem 8). They note that SpecDef’s compensation matrix could simply be combined with its partner, and argue that attacks cheaper than Theorem 8 are otherwise unlikely. Whether these limits hold on a real genomic model, and at what cost, has not been tested. If Theorem 8 were the cheapest informed attack, locking would still act as a cost barrier. A far simpler bypass exists. On every model we tested, a small change to how the compensation matrix is treated restores near-unlocked adaptation, and fusing the compensation matrix into its partner returns the original model, so the informed attack costs no more than fine-tuning an unlocked model.

We evaluate defence against two attacker tiers (Fig. 1; Methods). The *naive attacker* runs a standard fine-tuning pipeline on the released weights without knowledge of the lock, the adversary SpecDef was designed against. The *informed attacker* additionally knows the defence class and the public architecture and may modify tensors before fine-tuning. In this case, we evaluate the Theorem 8 construction and three interventions that need only an understanding of the defence, namely freezing the compensation matrix, fusing it into its partner, and canonicalising the network’s rescaling symmetries.

**Figure 1:**
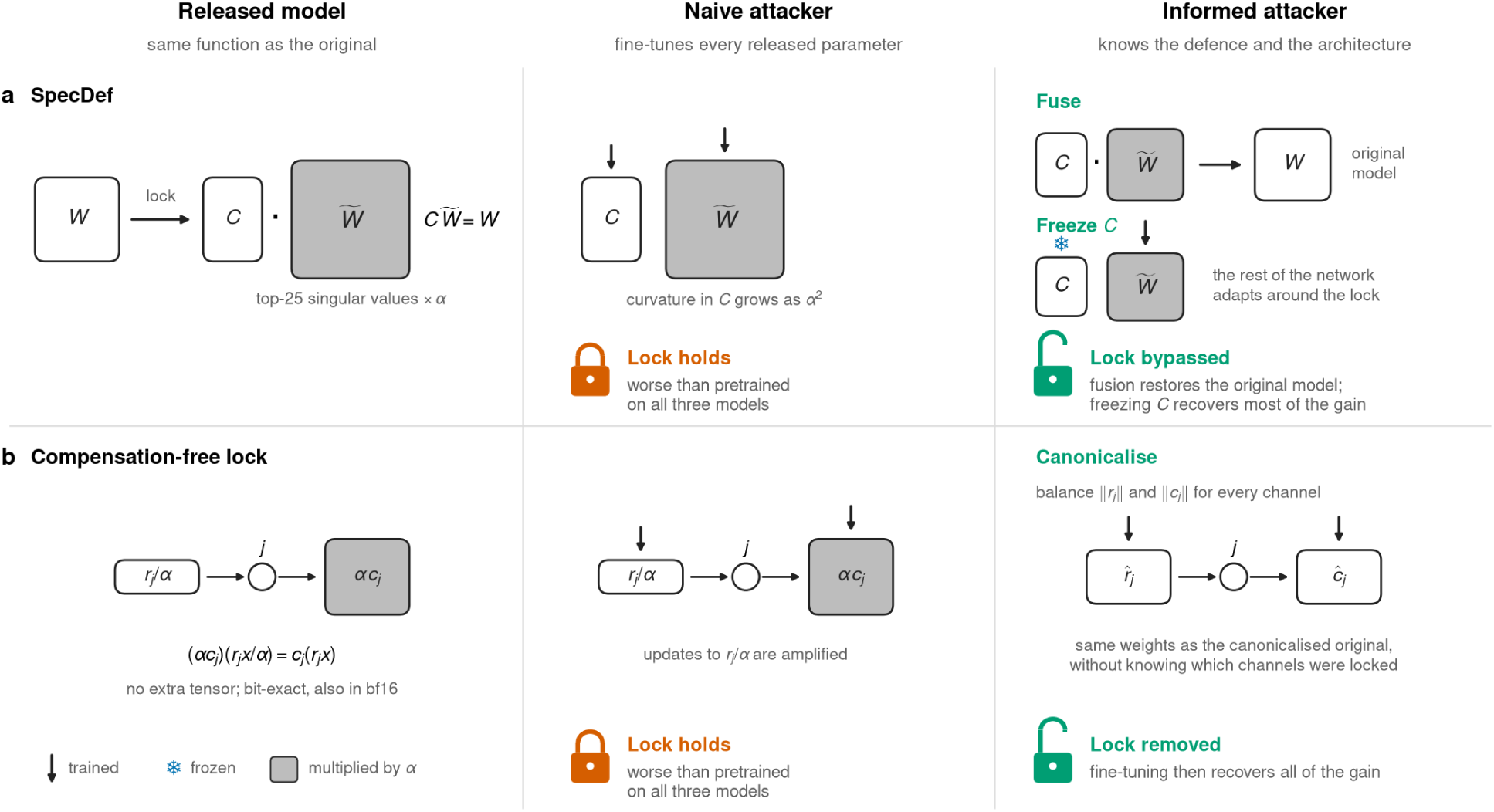
Two weight locks and two attackers. **(a)** SpecDef. **(b)** Compensation-free lock. Each row shows the lock as released (left), under the naive attacker (centre) and under the informed attacker (right), with the outcome on all three models. Arrows mark trained tensors, the snowflake a frozen tensor and grey tensors multiplied by *α*.

We applied SpecDef to Evo-2 7B and Evo-1-8k-base, and to the protein model ESM3-open (Hayes et al., 2025), from whose training data virus-specific sequences were likewise removed, and evaluated attack configurations spanning both tiers, under a fixed budget and explicitly controlled numerical precision. Our central finding is a separation by attacker tier. The lock defends against the naive attacker, whose full-parameter fine-tuning drives capability below the pretrained baseline on all three models, but the informed attacker removes it. A naive attacker who trains low-rank adapters meets the lock only if the adapters reach the tensor it amplifies. We then construct a compensation-free lock of the kind suggested previously (Rosati et al., 2026), folded into the network’s channel-rescaling symmetries. It defeats the naive attacker at no inference cost, but canonical rebalancing removes it. Two of these results go beyond the limits already established for this class of defence. The repair suggested by Rosati et al. (2026), a lock folded into the network’s own rescaling symmetries, is itself exactly removable, and provably so without knowledge of the lock (Proposition 1). And freezing all 32 targeted matrices, the strongest obstruction any lock placed on them could impose, costs the attacker almost nothing, so no defence confined to those matrices can hold, whatever its mechanism. Finally, we document how storage and compute precision determine whether resistance is observed at all.

## Results

### Fine-tuning recovers viral capability in an unlocked model

Consistent with previous reports on Evo-2 (Black et al., 2025; Wei et al., 2025), fine-tuning on the viral attack corpus for about one epoch (Methods) reduces held-out viral language-model loss significantly on both models (Fig. 2a; Supplementary Table 2). The gain is not memorization to genera seen in training and it remains substantial on held-out genomes whose genus is absent from the training split. The gain comes from the eukaryotic viral data, not from the extra training. Fine-tuning Evo-2 on the same amount of bacteriophage sequence, which it already saw in pretraining, leaves held-out loss at the pretrained value (1.833 against 1.834; Methods), in line with the phage control of Black et al. (2025).

**Figure 2:**
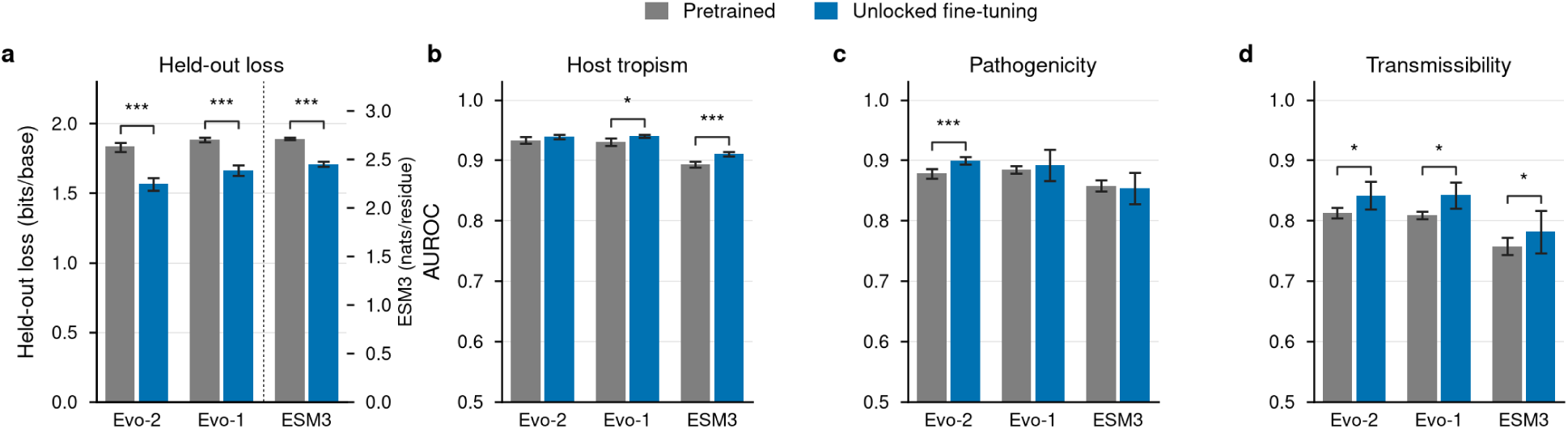
Viral adaptation after unlocked fine-tuning. **(a)** Held-out language-model loss of the pre-trained model and after unlocked fine-tuning. **(b–d)** Downstream AUROC for Host Tropism (**b**), Pathogenicity (**c**) and Transmissibility (**d**). Bars show means over training seeds and error bars 95% confidence intervals. Asterisks mark differences from the pretrained model (\**p* < 0.05, \*\**p* < 0.01, \*\*\**p* < 0.001; Supplementary Table 2).

After unlocked fine-tuning, the downstream classifiers (Methods) score higher on all three virological tasks on both models (Fig. 2b–d; Supplementary Table 2). The gains are significant for Transmissibility on both models, for Pathogenicity on Evo-2 and for Host Tropism on Evo-1. Because the HVUE tasks are proxies for virological properties and these gains are modest, held-out language modelling is our primary measure of adaptation.

### Spectral deformation preserves the released function

We applied SpecDef to the 32 write-side output projections of each model, inflating the top 25 singular values of each by *α* = 10^5^ on Evo-2 and 3 × 10^5^ on Evo-1 (Fig. 1a; Methods). At the main precision level, with the locked pair held in fp32, the released models reproduce the pretrained held-out loss to within 0.002 bits/base (Supplementary Fig. 1b). The lock therefore acts entirely on the optimisation problem, not on deployed behaviour. Served in bf16, however, the unfused lock degrades as *α* grows (Supplementary Fig. 1d), so a lock at the released strength must ship its locked layers in fp32.

### The lock defends against the naive attacker

Under standard fine-tuning of every released parameter, including *C*, the naive attacker fails to improve. On Evo-2 training destabilises, and on both models even the best checkpoint ends with a higher held-out loss and lower downstream performance on all three tasks than the pretrained model it started from (Fig. 3a,b; Supplementary Fig. 2 and Supplementary Table 3). On Evo-2 the naive attack fails at every learning rate, optimiser and budget we tried (Supplementary Table 4), consistent with the curvature with respect to *C* growing as *α*^2^ (Methods). Against the naive attacker, the lock therefore does more than deny the gain. It turns the attempt into a loss of capability.

**Figure 3:**
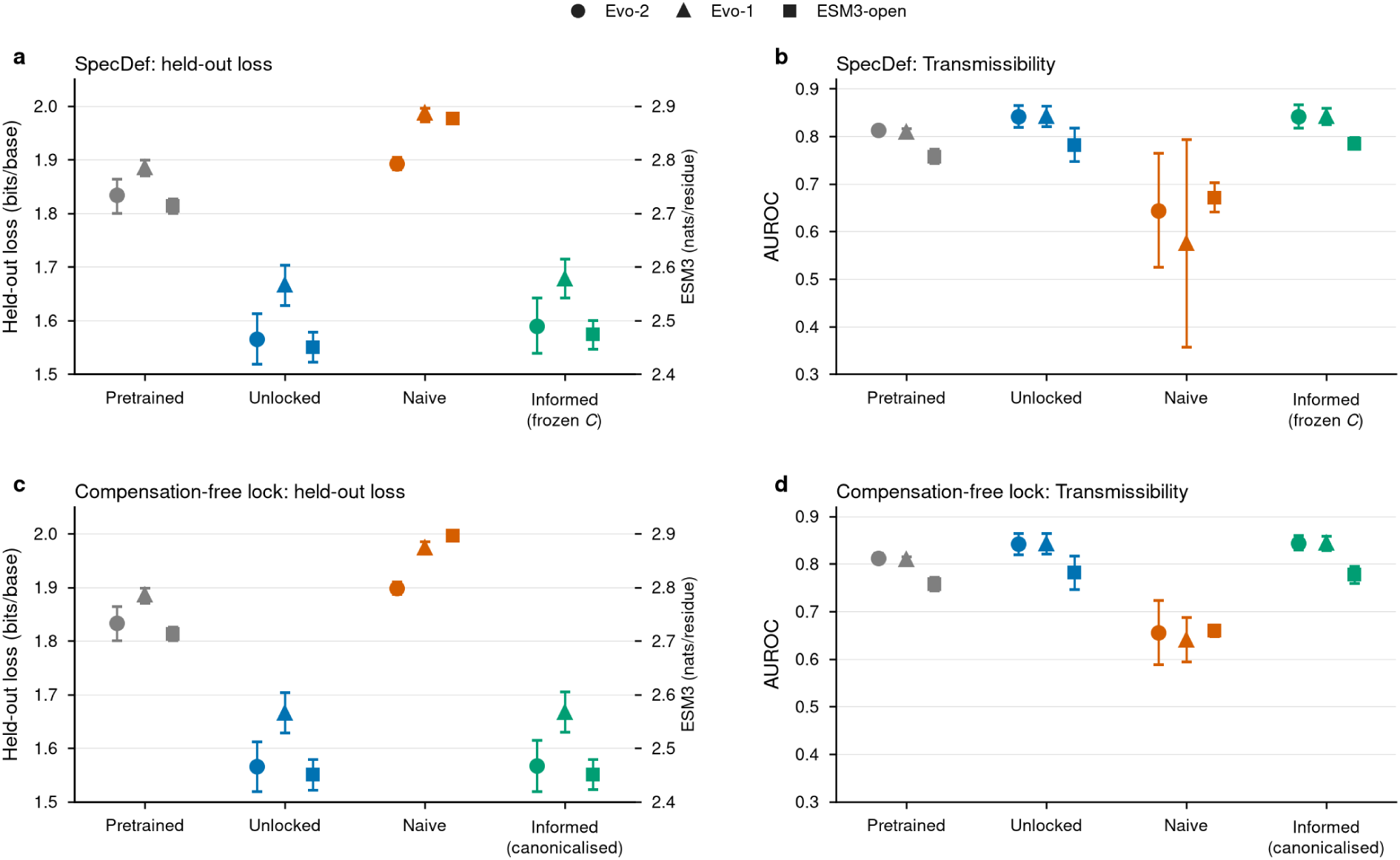
Held-out loss and Transmissibility AUROC for each attack. (a,b) SpecDef. **(c,d)** Compensation-free lock. Held-out language-model loss (**a,c**) and Transmissibility AUROC (**b,d**) at the best trained checkpoint, for the pretrained model, unlocked fine-tuning, the naive attacker and the informed attacker, who freezes *C* (**a,b**) or canonicalises the lock (**c,d**). Circles, Evo-2; triangles, Evo-1; squares, ESM3-open. Points show means and error bars 95% confidence intervals.

### The informed attacker removes the lock

The informed attacker can intervene and remove the locking parameterisation altogether. Because 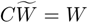 by construction (Eq. 1), an attacker who holds both factors computes their product and replaces the pair with the single fused matrix, as Rosati et al. (2026) anticipate. The released model then computes the original function. Its mean held-out loss matches the pretrained model to within 10^−5^ bits/base on both models. There is no subsequent attack to analyse, because the attacker is now fine-tuning the pretrained model (Supplementary Table 3).

A second intervention leaves the lock in place but makes *C* non-trainable. *C* keeps compensating in the forward pass, while the parameters whose curvature the lock inflates are removed from training. On Evo-2 this recovers 91% of the language-model gain (1.590 against 1.566 unlocked; difference +0.024, 95% CI +0.002 to +0.057), and all of it when the locked pair and its optimiser states are held at higher precision (1.564), consistent with the rounding effect described below. Downstream performance matches unlocked fine-tuning within the pre-registered equivalence margin on Host Tropism and Pathogenicity and is inconclusive on Transmissibility (Fig. 3a,b). The same pattern holds on Evo-1, where the pre-registered equivalence criteria for held-out loss and Transmissibility are met (Supplementary Table 3).

Both interventions are far cheaper than the Theorem 8 chain, which on Evo-1 costs up to 2.6× the per-step time of unlocked fine-tuning (Supplementary Table 1).

Freezing *C* does not undo the lock. With *C* frozen, the 25 inflated directions of each locked matrix move by 1.2×10^−8^ of ||*W*|| over training, against 6.4×10^−3^ in an unlocked run, while their orthogonal complement moves by 2.1 × 10^−2^ against 4.0 × 10^−2^ (Fig. 4a). The lock still holds where it was placed and the rest of the network adapts around it.

**Figure 4:**
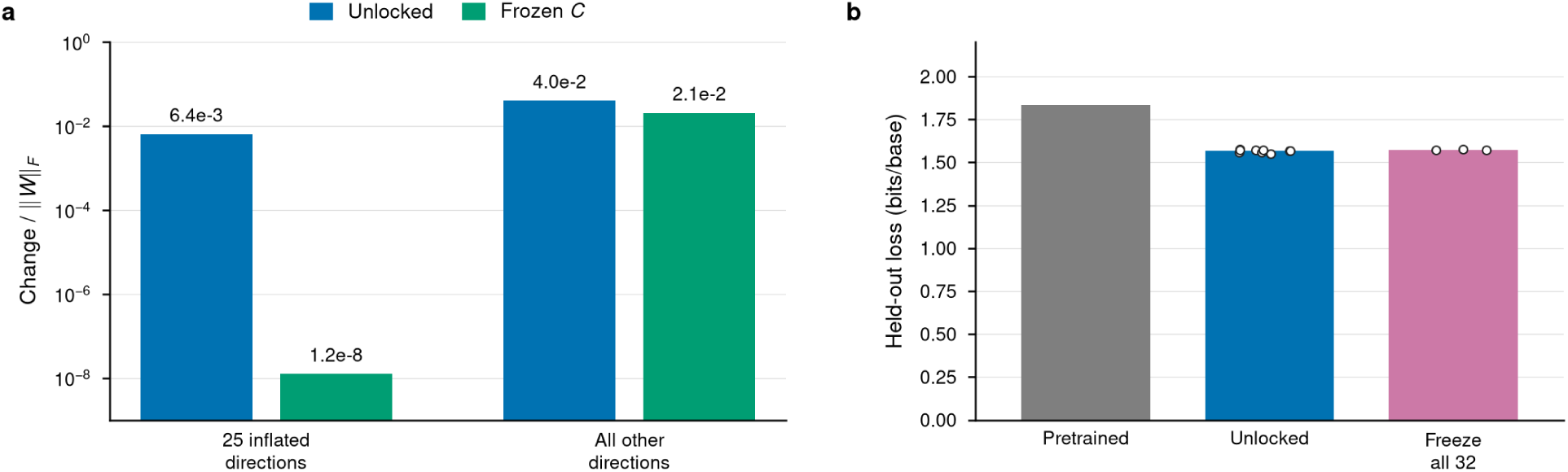
Weight change in the locked matrices and freezing of all targeted matrices. Evo-2, SpecDef at *α* = 10^5^. **(a)** Change in each locked matrix during fine-tuning, as a fraction of its norm, along the 25 inflated directions and along all other directions, with *C* frozen and in unlocked fine-tuning (log scale). **(b)** Held-out loss (bits per base) after fine-tuning with all 32 targeted matrices frozen, compared with unlocked fine-tuning and the pretrained model. Bars show means; points in **b** show individual training seeds.

### Freezing the targeted matrices

Freezing all 32 targeted matrices outright is the strongest obstruction any lock placed on them could impose on an attacker who leaves them untrained. It still recovers 97% of the language-model gain, not significantly less than unlocked fine-tuning, and with no significant downstream cost (Fig. 4b and Supplementary Table 3). No defence confined to these matrices can therefore produce a model-level barrier against such an attacker, whatever its mechanism.

### Compensation-free lock

SpecDef hands the attacker a separate, clearly labelled compensation tensor that can be easily identified and bypassed. Locks built from per-channel rescaling have been argued to be reversible in principle, on the grounds that an attacker can inspect per-layer norms and rebalance them (Sakamoto et al., 2026), but the argument has not been tested on a model of this size. We therefore constructed the strongest version of the idea available to a defender by producing a lock with no additional tensor, no inference cost, and no object to freeze or fuse.

The lock divides the row that produces a hidden channel by *α* and multiplies the column that consumes it by *α*, which leaves the function unchanged while separating the two factors in norm by *α*^2^ (Fig. 1b; Methods). The construction is bit-exact in bf16 at every strength tested (2^17^ to 2^56^) on both models.

The naive attacker is defeated again, in both language modelling and downstream performance. On Evo-2 at *α* = 2^32^ the best checkpoints reach 1.898 bits/base on average against 1.834 pretrained, and all three tasks fall below pretrained. Evo-1 at *α* = 2^31^ shows the same (1.974 against 1.885; Fig. 3c,d and Supplementary Table 3). The informed attacker canonicalises and bypasses the defensive mechanism, moving every channel to the point where its producer row and consumer column have equal norms, a standard rebalancing of rescaling symmetries (Stock et al., 2019).

Canonicalisation takes about 50 lines of code and one GPU-minute, and needs only the public channel layout, not the original weights, the modified channels, *α* or knowledge that the model was locked. The canonicalised locked model is bit-identical to the canonicalised original, every tensor, at every strength on all models, and fine-tuning it recovers unlocked behaviour. The same holds for the strongest version of the construction, which applies every valid per-channel rescaling of Evo-2 at once (Methods).

### Parameter-efficient fine-tuning

A naive attacker may instead train low-rank adapters (LoRA) on a chosen set of layers. Such an attacker meets a lock only if an adapter sits on the tensor whose updates the lock amplifies, which is *C* for SpecDef and the producer rows for the compensation-free lock. An attacker unaware of the lock has no reason to exclude these tensors, and adapting every linear layer, as QLoRA recommends (Dettmers et al., 2023), includes them. Both locks then hold on Evo-2, consistent with the divergence reported for SpecDef under LoRA (Rosati et al., 2026). Narrower conventions can miss these tensors without any knowledge of the lock. Adapters on the input projections alone leave SpecDef invisible, although the compensation-free lock still holds, and adapters on the output projections alone bypass both. Both also hold on Evo-1 (Methods).

### The separation holds on a protein language model

We repeated the core experiments on ESM3-open, using the proteins of the same viral genomes (Methods). Unlocked fine-tuning lowers held-out masked-token loss significantly (Supplementary Table 2) and improves Host Tropism and Transmissibility (Fig. 2). We locked its 48 attention output projections with SpecDef (*α* = 10^5^) or the compensation-free lock (2^32^) and both releases preserve its function. Both locks defeat the naive attacker. The informed attacker again removes them. Freezing *C* or all 48 locked pairs recovers about 91% of the gain (not equivalent to unlocked within the pre-registered margin, as for Evo-2 in fp32), fusion returns the original model, and canonicalisation recovers all of the gain (Fig. 3; Supplementary Table 5).

### Measured resistance depends on numerical precision

Whether the lock appeared to work depended repeatedly on how numbers were stored. A stored number can only change in steps of a minimum size, which grows with its magnitude and shrinks with the precision of the format. In fp32, a weight of size 10^4^ changes in steps of about 10^−3^, so an update of 10^−5^ is lost.

#### The suppression inside the inflated matrices is a rounding effect

When only the 32 inflated matrices 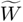 are trained, a lock stored in fp32 learns about a third as much as an unlocked control, whereas stored in f64 it learns like the control (Supplementary Fig. 1a). The curvature with respect to 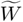 does not change with *α* (Methods), so these updates are lost to the rounding of the inflated weights, not to curvature. This is related to the numerically ineffective regime described previously (Rosati et al., 2026), and it is consistent with freezing *C* recovering slightly less than the full gain in fp32.

#### A broken model can look like a protected one

A lock that stops preserving the original function damages the released model, and a damaged model also fails to improve under fine-tuning. Storing *C* in bf16 breaks the identity 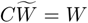 before training starts (Supplementary Fig. 1b), and so does loading the released model with the 4-bit (Dettmers et al., 2023) or 8-bit (Dettmers et al., 2022a) quantisation of memory-saving fine-tuning recipes, under which both locked Evo-2 models lose their function before any training, as noted previously (Rosati et al., 2026). Rounding to bf16 breaks SpecDef but leaves the compensation-free lock exact. An evaluation that only asks whether the attacker improved cannot tell such a model from a working defence, since an attacker who quantises the locked model starts from a broken one, and neither lock can be served quantised.

Storage precision also limits the attacker. With weights stored in bf16, the default, roughly 83% of weights never change and even unlocked fine-tuning adapts about half as far as with fp32 storage (Supplementary Fig. 1c), which is why all headline results store weights in fp32.

## Discussion

Both locks we evaluate obstruct an attacker who does not know they are there, and neither obstructs one who does. The barrier is one of knowledge, not compute. An informed attacker removes or circumvents both with a simple change made before fine-tuning, at no more than the cost of fine-tuning an unlocked model, and some low-rank adapter conventions bypass SpecDef without any knowledge of the lock at all. Weight locking does defeat naive full-parameter fine-tuning on two Evo-family models and ESM3-open, so its value as a governance mechanism is real but inherently limited, because it depends on attackers not knowing a bypass that is now published.

These findings instantiate the limit proved by Rosati et al. (2026). Their constructive bypass restores trainability of the locked layers at linear cost in model size, and we find that it runs at seven-billion-parameter scale. But the attacker does not need to restore trainability of the locked layers. They can simply route around them, which freezing all 32 targeted matrices shows costs almost nothing, or move to coordinates in which the same function is easy to optimise.

For locks built from an explicit compensation matrix or from per-channel rescaling, the limitation is structural. Embedding the lock in the network’s intrinsic rescaling symmetries yields a construction strictly better for the defender, with no extra tensor, no inference cost and bit-exact release, and it remains exactly removable by canonicalisation. The result joins a broader pattern in which training-time safeguards for open-weight models fail under systematic re-evaluation (Qi et al., 2025; Tamirisa et al., 2025; Rosati et al., 2024).

Our study has limitations. We tested two Evo-family models and one transformer protein model. For two of the three we ran the core attacks at one lock strength and one training recipe because of compute cost. The adaptation we seek to prevent is characterised on a single corpus of 544 viral genomes.

In conclusion, we confirm that excluding eukaryotic viral genomes from training is not a sufficient safeguard on its own, because the pretrained models acquire much of this capability within a few GPU-hours of fine-tuning. Weight locking adds protection only against the naive attacker, and the locks are also indiscriminate, since they obstruct legitimate adaptation, such as variant interpretation or regulatory-element prediction, as much as misuse. The unlocked-attacker results reported here provide a baseline against which to test new safeguards on Evo-family models and ESM3-open. We believe the field should move towards methods that remove the capability itself and make it hard to relearn, such as targeted unlearning by gradient difference (Liu et al., 2022) or representation misdirection (Li et al., 2024), guided by causal localisation (Meng et al., 2022), and representation noising (Rosati et al., 2024).

## Methods

### Models

We used Evo-1-8k-base (Nguyen et al., 2024) and Evo-2 7B (Brixi et al., 2026). Evo-1-8k-base has 7 billion parameters in 32 blocks (29 Hyena operators and 3 multi-head attention layers), a hidden dimension of 4,096 and a single-nucleotide tokenizer. Evo-2 7B has 32 blocks (27 Hyena operators and 5 multi-head attention layers), a hidden dimension of 4,096 and a byte-level single-nucleotide tokenizer. ESM3-open (`esm3-sm-open-v1`) (Hayes et al., 2025) is a 1.4-billion-parameter transformer with 48 blocks and a hidden dimension of 1,536, trained on protein sequence, structure and function with sequences unique to viruses removed from its training data and we use its sequence track only.

### Data

The attack corpus consists of 910 NCBI Virus (Hatcher et al., 2017) assemblies of eukaryotic viruses, including influenza A virus and SARS-CoV-2, split by accession into 544 training and 366 held-out genomes. Held-out language modelling uses 256 windows of 1,024 nucleotides, drawn once with a fixed seed by choosing a held-out genome uniformly at random (with replacement) and a start position uniformly within it. A second set of 128 windows, drawn the same way from the 58 held-out genomes whose genus is absent from the training split, measures the gain on genus-disjoint sequence. Downstream evaluation uses the Host Tropism, Pathogenicity and Transmissibility tasks of HVUE (Dutta et al., 2026), with the version 2 hard splits. The bacteriophage control uses 544 fragments of RefSeq (O’Leary et al., 2016) phage genomes, each from a different genome and cut to the length of one training genome, so that genome count, length distribution and total length (24.47 Mb) match the training split. For ESM3-open, the corpus is the annotated protein translations of the same genomes (16,279 training and 9,537 held-out proteins, after removing duplicates and held-out proteins identical to a training protein), and each HVUE fragment is translated to the longest stop-free stretch over its six reading frames, which keeps every example, label and split.

### Threat model

The *naive attacker* holds the released weights and a viral corpus and fine-tunes the released model with a standard recipe, training either all released parameters, including any compensation matrices, or low-rank adapters on a chosen set of layers, without inspecting or modifying the weights. The *informed attacker* also knows the defence and the public architecture, and may freeze, fuse or rebalance tensors before fine-tuning. Neither has access to the original weights, the modified directions or channels, or the lock strength *α*. A defence *holds* when the attacker’s best checkpoint is not significantly better than the pretrained model on held-out language modelling (the 95% genome-bootstrap interval of the difference does not lie entirely below zero).

### Fine-tuning

Full-parameter runs use AdamW (Loshchilov and Hutter, 2019) at a learning rate of 10^−5^ with 10% linear warm-up, no weight decay, gradient clipping at 1.0 and precision level L1. On the Evo models the headline recipe is 6,000 steps of 4,096 tokens (24.6 M tokens, about one epoch) with 8-bit optimiser states (Dettmers et al., 2022b). The training seeds and precision level of each arm are given with its result in the Supplementary Tables.

On Evo-2, multiple learning rates were tested, gradient clipping, optimiser and budget one at a time, each setting paired with an unlocked run trained the same way (Supplementary Table 4). Adafactor (Shazeer and Stern, 2018) and SGD with momentum 0.9 were run at the best unlocked learning rate from a grid and at a tenfold lower one, and the 24,000-step runs (about four epochs) use fp32 optimiser states and a 600-step warm-up. The phage control applies the unlocked recipe to the phage corpus (three seeds), selecting checkpoints on the eukaryotic monitor windows. On Evo-1, the equivalence tests (frozen *C*, canonicalisation) and the compensation-free lock use fp32 optimiser states and are compared with an unlocked run trained the same way.

ESM3-open is fine-tuned with a masked-token objective (15% of residues masked) and fp32 optimiser states for 3,000 steps of 16 crops of up to 512 residues, the Evo token budget. Each example draws a genome uniformly and then one of its proteins in proportion to length, as for the Evo windows. A first run that drew proteins uniformly put 87% of training on three families of large DNA viruses and was repeated as a pre-registered deviation, with the same conclusions about the locks. SpecDef (*α* = 10^5^, *k*_lock_ = 25) is applied to the 48 attention output projections, and the compensation-free lock (2^32^, 25 channels per block) to the value rows of each attention input projection and the matching columns of its output projection.

Low-rank attacks train rank-16 LoRA adapters (Hu et al., 2022) (scaling *α*_LoRA_*/r* = 2, unrelated to the lock strength) with every base weight frozen, at 10^−4^ with fp32 optimiser states, 10% warm-up and the headline budget. Adapters go on every linear layer, including both 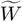 and *C* of each SpecDef pair, on the input projections only (Hyena `projections`, attention `Wqkv`) or on the output projections only (`out_filter_dense`, `out_proj`, with *C* unadapted). A fourth list reproduces a library’s all-linear setting, which skips Evo-2’s Hyena input projections because they are not `torch.nn.Linear`. On Evo-1 only the every-linear-layer list was run. A separate attack trains the task classifiers directly on the released Evo-2 lock, with rank-16 adapters on 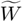 alone or on both 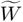 and *C* inside the 27 locked Hyena output projections, the five locked attention projections frozen and no language-model fine-tuning.

### Numerical precision

Results are reported at one of three precision levels. L0 stores weights in bf16, as the models load by default. L1 stores weights in fp32, computes in bf16 and holds each locked pair in fp32; all headline results use L1. L2 is L1 with the locked pair held in f64 and its optimiser states in full precision. The precision of the attacker’s weights, the optimiser states and the locked tensors is recorded for every run. In the experiment of Supplementary Fig. 1a, only the 32 inflated matrices 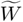 are trained, with *C* and all other parameters frozen, with AdamW with 8-bit states and three seeds, and learning is measured as the fall in perplexity on 64 fixed monitor windows.

### Held-out language modelling

Language-modelling endpoints are reported in bits per base over the 256 held-out windows. A run’s best checkpoint is the trained checkpoint with the lowest loss on 16 fixed monitor windows from the held-out genomes. It is then scored on the 256 held-out windows. Confidence intervals are obtained by bootstrap resampling of genomes (10,000 resamples, 95% percentile intervals, training seeds resampled within each arm). For ESM3-open, held-out loss is the masked-token cross-entropy (nats per masked residue) on all 9,537 held-out proteins with fixed masks, pooled within each of the 364 held-out genomes and then averaged over genomes, so that each genome counts equally as in the Evo windows. Checkpoints are selected on 64 held-out proteins drawn the same way.

### Downstream classifiers

Downstream performance is measured by fine-tuning each checkpoint on each HVUE task. For every checkpoint and task, rank-16 LoRA adapters (Hu et al., 2022) are trained together with a classification head (batch normalisation and a linear layer on mean-pooled hidden states), with the checkpoint’s own weights frozen. Inputs are up to 1,000 nucleotides. Classifiers use AdamW (learning rate 10^−4^, no weight decay, effective batch 32) for up to 4,000 steps; validation AUROC is computed every 500 steps and the best validation checkpoint is scored on the test split. Test AUROC is reported on the examples without near-duplicates in the training split. For ESM3-open the classifiers follow the same protocol on the translated tasks, with adapters on the attention and feed-forward linear maps of the 48 blocks and mean-pooling over residue positions only.

### Spectral deformation

For a target matrix with thin SVD *W* = *U*Σ*V*^T^, SpecDef releases the pair

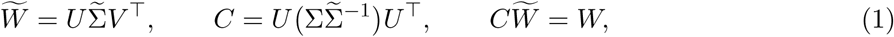

where the top *k*_lock_ = 25 singular values of 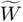 are inflated by *α*. SpecDef was applied to the 32 projections that write each block’s output, the `out_filter_dense` matrices of the Hyena blocks and the `out_proj` matrices of the attention blocks (4096 × 4096 each; about 537 M parameters in total), at *α* = 10^5^ on Evo-2 and Evo-1. Forward preservation was verified at deployment precision before fine-tuning.

### Informed attacks on SpecDef

*Freezing* makes every compensation matrix *C* non-trainable and trains all other parameters with the headline recipe. *Fusion* computes each product 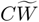 in f64 and replaces the pair with the resulting matrix before training. We report its reconstruction error at deployment precision. The *SVD-chain attack* implements Theorem 8 of Rosati et al. (2026) on Evo-1. Each 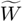 is replaced by *k* ∈ {2, 3, 5} trainable factors initialised from its *k*-th matrix root, reducing the per-factor spectral norm from 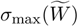 to 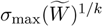, while *C* is held fixed. Per-step time was measured on Evo-1 with AdamW with 8-bit states at *α* = 3 × 10^5^, on one GH200 per run, as the mean over updates 101–500 of a 501-update run, for unlocked fine-tuning, the naive attack, freezing and fusion at precision level L1, and for the SVD chain with float64 factors whose optimiser states are held in host memory and with the network stored in bf16 (level L0), because the chain with float64 factors ran out of GPU memory at L1. Each configuration is reported relative to an unlocked run at the same precision level.

### Freezing the targeted matrices

All 32 targeted matrices, and *C* where present, are frozen and the remaining parameters are trained with the headline recipe.

### Parameter movement and curvature

For each locked matrix, the change in weights over training is projected onto the *k*_lock_ inflated left-singular (output-side) directions and onto their orthogonal complement. Each projection is reported as a fraction of ||W||_F_ averaged over the 32 matrices and three seeds. The top Hessian eigenvalue of the loss with respect to the *C* block, the 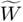 block and both jointly was estimated on Evo-2 by power iteration on finite-difference Hessian–vector products, with fp32 compute and the locked pair in f64.

### Compensation-free lock

In each of the 32 output projections we select the 25 input channels with the largest column norm. For each selected channel, the consuming column is multiplied by *α* = 2*^E^* and the producing row of the upstream tensor is divided by *α*. In matrix form, for adjacent maps *W*_1_ (producer) and *W*_2_ (consumer) and a diagonal matrix *D* holding *α*^−1^ on the selected channels and 1 elsewhere,

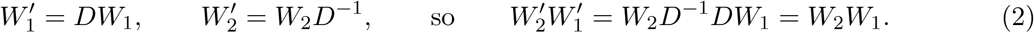

This preserves the model’s function only if every operation between producer and consumer scales each channel independently and linearly. In attention blocks the producer is the value projection and the consumer is `out_proj`. The attention weights depend only on queries and keys and mix positions rather than channels, so the rescaling is exact. In Hyena blocks the producer is the input-projection row that generates the channel and the consumer is `out_filter_dense`. The short and long convolutions act on each channel separately, and elementwise gating preserves the rescaling provided the channel enters only one of the multiplied branches. A normalisation layer, a softmax over channels or a residual connection between producer and consumer would break the rescaling. Forward preservation was verified by exact tensor comparison at bf16, fp32 and f64 at every *α* tested.

### Canonicalisation

For a channel *i* with producer row *r_i_* and consumer column *c_i_*, multiplying *r_i_* by any *s* > 0 and dividing *c_i_* by the same *s* leaves the model unchanged (Eq. 2). Canonicalisation fixes this free choice by picking, for every channel, the value of *s* that makes the row and the column equally large,

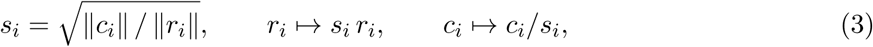

after which both have norm 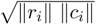. In practice *s_i_* is rounded to a power of two, 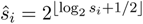. We apply this to every channel of every producer–consumer pair in which rescaling leaves the model unchanged. Rescaling symmetries (Dinh et al., 2017) and their balancing (Stock et al., 2019) are established.

For the strongest compensation-free lock on Evo-2 we also rescale, in 25 channels per pair at *α* = 2^32^, the normalisation weights against the linear maps they feed and the hidden channels of each MLP against its output map. The release remains bit-exact. These pairs share tensors, so Proposition 1 does not apply directly. Canonicalisation therefore balances all pairs in a fixed order and repeats until a full sweep changes nothing, and the canonicalised locked and original models then agree to within 10^−12^ without rounding and bit for bit with rounding. On ESM3-open the rescaled pairs are disjoint, so a single pass suffices.

**Proposition 1** (Canonicalisation ignores the lock). For channels whose producer and consumer belong to no other rescaled pair, canonicalisation (Eq. 3) gives the same weights for the original model and for any rescaled version of it, including the locked model.

*Proof.* Rescaling channel *i* by *d >* 0 multiplies ||*r_i_*|| by *d* and divides ||*c_i_*|| by *d*. It changes neither the directions of *r_i_* and *c_i_* nor the product ||*r_i_*|| ||*c_i_*||. After canonicalisation the row has the direction of *r_i_* and norm 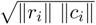, and the column likewise, so the result does not depend on *d*.

## Data availability

The HVUE benchmark is publicly available (Dutta et al., 2026). The attack corpus accession lists and the phage and protein corpora definitions are available at https://github.com/Georgakopoulos-Soares-lab/glm-locking. Evo-1-8k-base and Evo-2 weights are from their original authors and ESM3-open weights from EvolutionaryScale; we redistribute no model weights.

## Code availability

Locking and canonicalisation scripts, fine-tuning configurations, the evaluation pipeline and the audit record are available at https://github.com/Georgakopoulos-Soares-lab/glm-locking.

## Dual-Use Statement

This work evaluates a safeguard intended to limit the misuse of open-weight biological models. The capability we measure is adaptation on task proxies, namely held-out viral language modelling and three HVUE classification tasks; it does not demonstrate the ability to design pathogens, and we report no generated sequences or design protocols. The attack corpus consists of public NCBI Virus and RefSeq records, the downstream tasks use the public HVUE benchmark, and no wet-laboratory work was performed. The bypasses we describe defeat a defence that is not deployed at scale. We judged that publishing them is beneficial, because otherwise the defence could be adopted on the strength of results that do not hold against an informed attacker. We release code, configurations and the audit record, but no fine-tuned attack checkpoints.

## Author Contributions

I.M., I.G.-S. and A.K. conceived the study. A.K. performed the experiments and analysed the data with the help of A.V. A.K., C.S.C., I.M. and I.G.-S. wrote the manuscript. I.M. and I.G.-S. jointly supervised the work. All authors reviewed the manuscript.

## Supplementary Information

**Supplementary Fig. 1:**
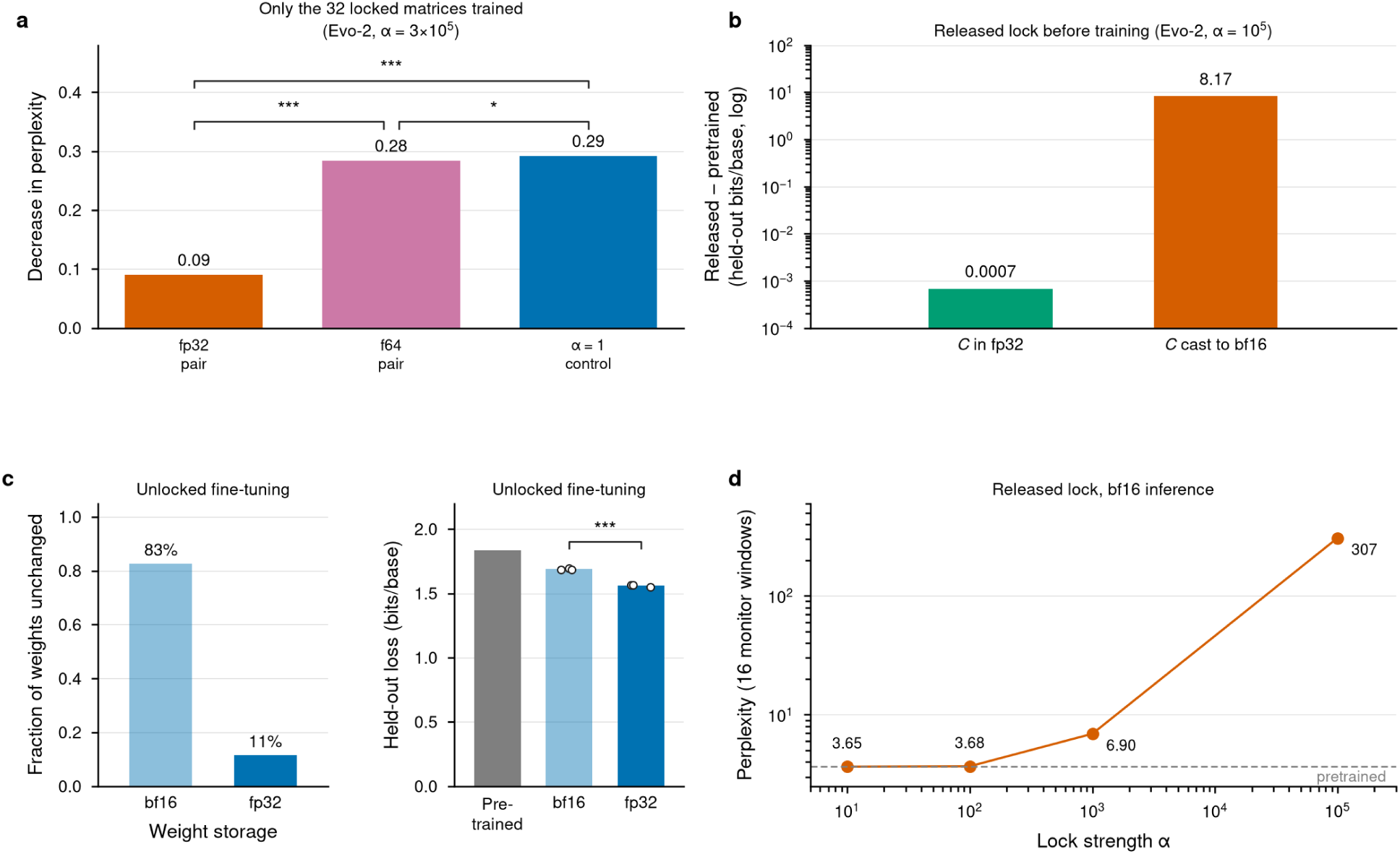
Numerical precision. Evo-2. **(a)** Fall in perplexity when only the 32 inflated matrices 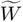 are trained (*C* and all other parameters frozen), for a SpecDef lock (*α* = 3 × 10^5^) stored in fp32 or f64 and for an unlocked control. **(b)** Difference in held-out loss between the released SpecDef lock (*α* = 10^5^) and the pretrained model, with *C* stored in fp32 or cast to bf16 (log scale). **(c)** Fraction of weights that never change, and held-out loss, in unlocked fine-tuning with weights stored in bf16 or fp32. **(d)** Perplexity of the released SpecDef lock under bf16 inference against lock strength *α*. Bars show means and points individual training seeds. Asterisks mark significant differences (\**p* < 0.05, \*\**p* < 0.01, \*\*\**p* < 0.001; Welch’s *t*-test in **a**, genome-level bootstrap in **c**).

**Supplementary Fig. 2:**
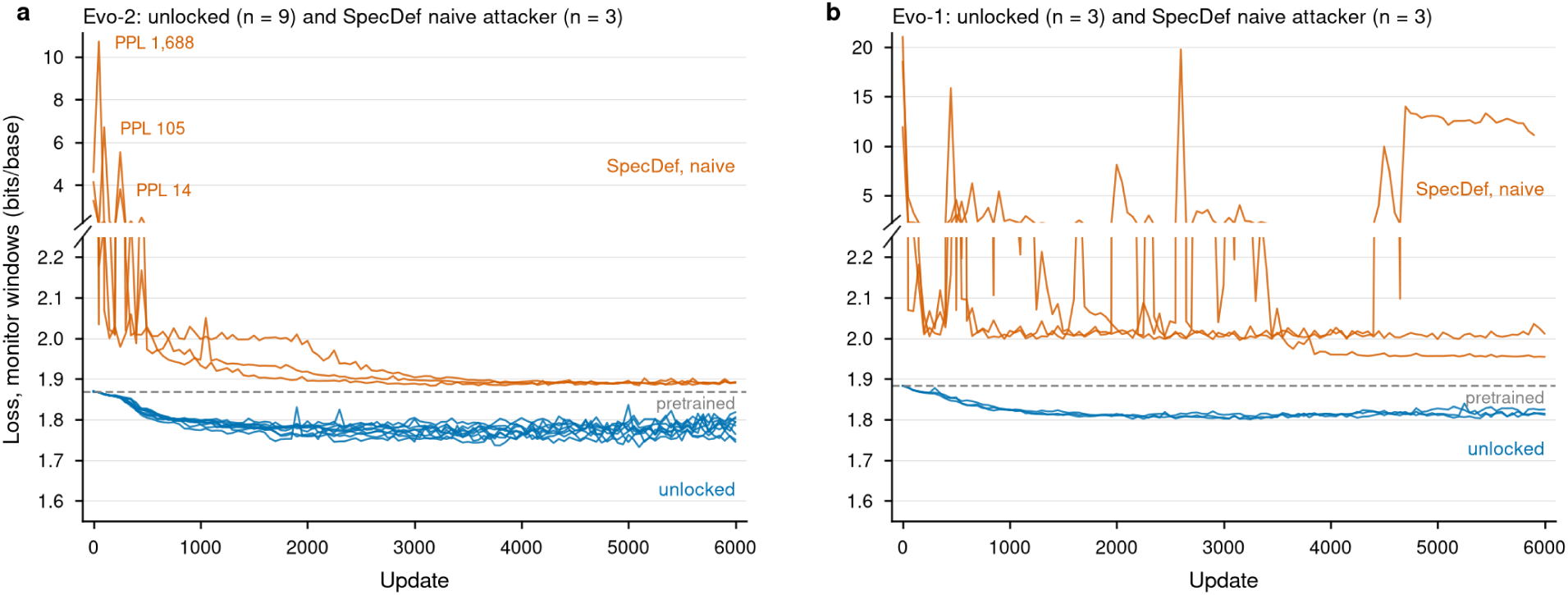
Training curves. Loss on the 16 monitor windows (bits per base) against training step, for unlocked fine-tuning and the naive attack on SpecDef, on Evo-2 and Evo-1. Each line is one training seed.

**Supplementary Table 1:**
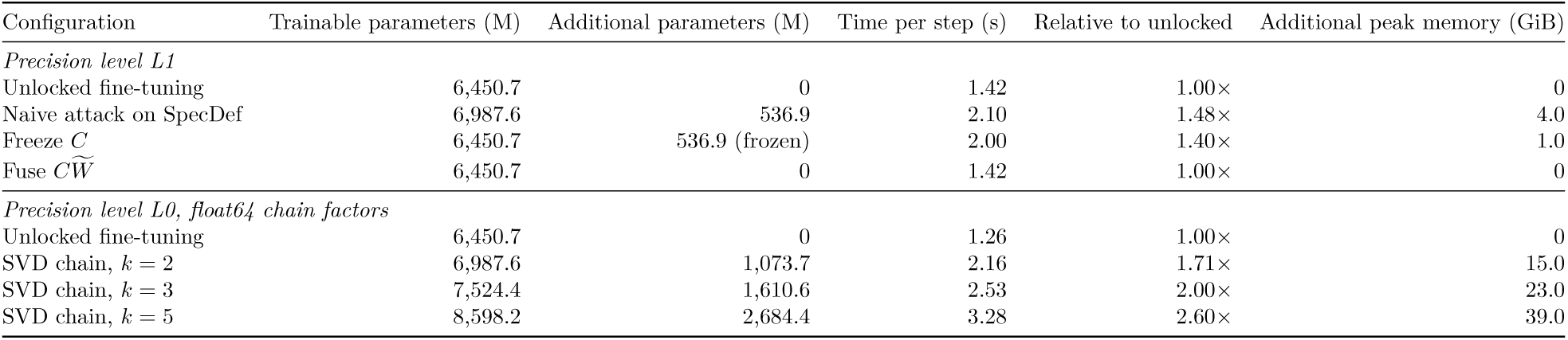
Time, parameters and memory of each attack configuration. Evo-1, AdamW with 8-bit states, *α* = 3 × 10^5^, one GPU per run. Time per step is averaged over updates 101–500, and parameters and peak GPU memory are given relative to unlocked fine-tuning at the same precision level (Methods).

| Configuration | Trainable parameters (M) | Additional parameters (M) | Time per step (s) | Relative to unlocked | Additional peak memory (GiB) |
|---|---|---|---|---|---|
| *Precision level L1* |  |  |  |  |  |
| Unlocked fine-tuning | 6,450.7 | 0 | 1.42 | 1.00× | 0 |
| Naive attack on SpecDef | 6,987.6 | 536.9 | 2.10 | 1.48× | 4.0 |
| Freeze *C* | 6,450.7 | 536.9 (frozen) | 2.00 | 1.40× | 1.0 |
| Fuse *CW̃* | 6,450.7 | 0 | 1.42 | 1.00× | 0 |
| *Precision level L0, float64 chain factors* |  |  |  |  |  |
| Unlocked fine-tuning | 6,450.7 | 0 | 1.26 | 1.00× | 0 |
| SVD chain, *k* = 2 | 6,987.6 | 1,073.7 | 2.16 | 1.71× | 15.0 |
| SVD chain, *k* = 3 | 7,524.4 | 1,610.6 | 2.53 | 2.00× | 23.0 |
| SVD chain, *k* = 5 | 8,598.2 | 2,684.4 | 3.28 | 2.60× | 39.0 |

**Supplementary Table 2:** Difference between unlocked fine-tuning and the pretrained model. Held-out loss (bits per base, or nats per masked residue for ESM3-open) and downstream AUROC. Unlocked values are means over training seeds. Differences, 95% confidence intervals and *p* values come from a genome bootstrap for held-out loss and, for AUROC, from a linear mixed model with training seed and classifier seed as crossed random effects (Evo-2 Pathogenicity and Transmissibility, nine training seeds; ESM3-open) or a Welch test (Evo-2 Host Tropism, nine training seeds; Evo-1), Holm-corrected across the three tasks of each model.

| Model | Measure | Pretrained | Unlocked | Difference [95% CI] | *p* |
|---|---|---|---|---|---|
| Evo-2 | Held-out loss (bits/base) | 1.834 | 1.566 | −0.268 [−0.317, −0.220] | < 0.001 |
| Evo-2 | Host Tropism AUROC | 0.933 | 0.939 | +0.005 [−0.001, +0.011] | 0.080 |
| Evo-2 | Pathogenicity AUROC | 0.878 | 0.899 | +0.022 [+0.011, +0.033] | < 0.001 |
| Evo-2 | Transmissibility AUROC | 0.813 | 0.842 | +0.028 [+0.008, +0.049] | 0.015 |
| Evo-1 | Held-out loss (bits/base) | 1.885 | 1.665 | −0.220 [−0.258, −0.184] | < 0.001 |
| Evo-1 | Host Tropism AUROC | 0.930 | 0.940 | +0.010 [+0.003, +0.016] | 0.031 |
| Evo-1 | Pathogenicity AUROC | 0.884 | 0.892 | +0.008 [−0.014, +0.030] | 0.314 |
| Evo-1 | Transmissibility AUROC | 0.809 | 0.842 | +0.033 [+0.015, +0.051] | 0.031 |
| ESM3-open | Held-out loss (nats/residue) | 2.713 | 2.450 | −0.263 [−0.290, −0.237] | < 0.001 |
| ESM3-open | Host Tropism AUROC | 0.893 | 0.910 | +0.016 [+0.013, +0.020] | < 0.001 |
| ESM3-open | Pathogenicity AUROC | 0.858 | 0.854 | −0.004 [−0.021, +0.013] | 0.635 |
| ESM3-open | Transmissibility AUROC | 0.758 | 0.782 | +0.024 [+0.007, +0.042] | 0.014 |

**Supplementary Table 3:**
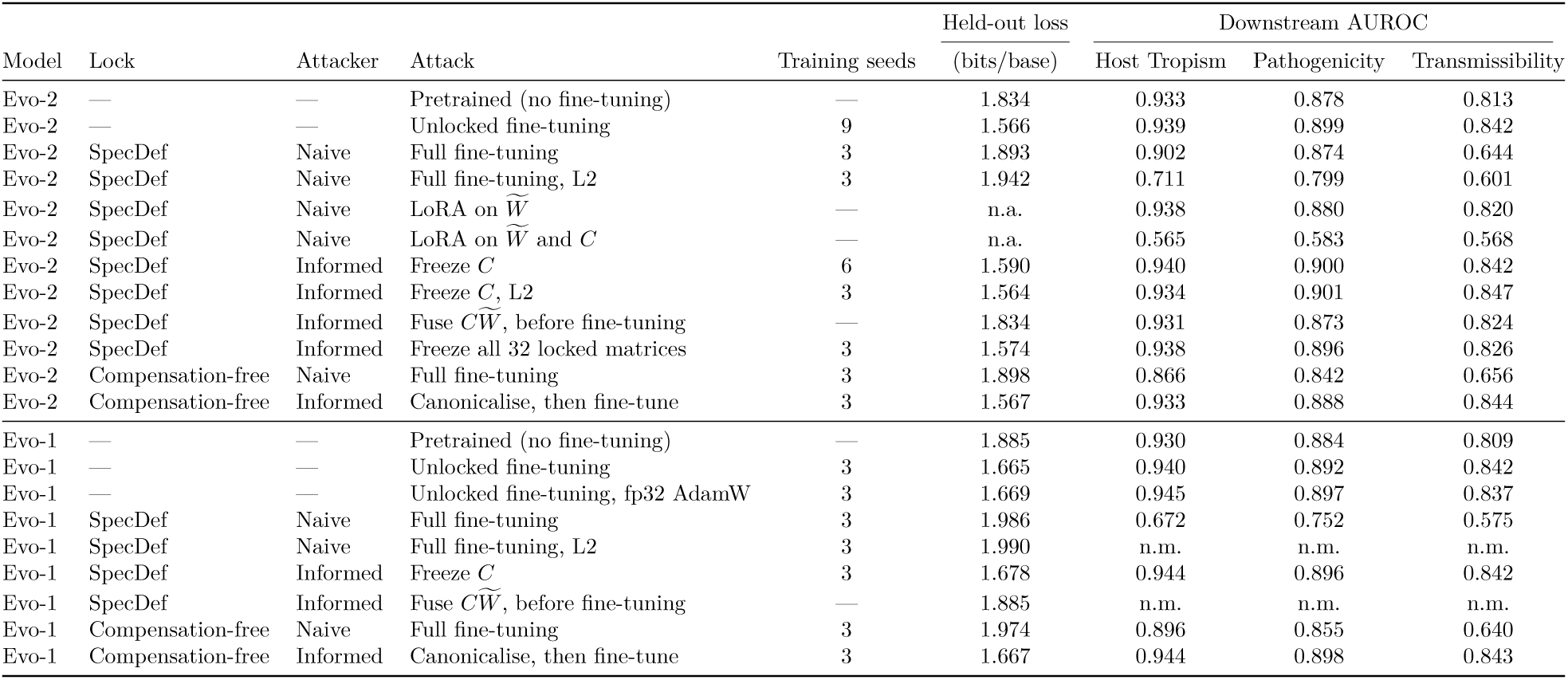
Held-out loss and downstream AUROC for each attack. Evo-2 and Evo-1, at the best trained checkpoint. Values are means over training seeds, with AUROC first averaged over classifier seeds. SpecDef at *α* = 10^5^ (Evo-2) and 3 × 10^5^ (Evo-1), compensation-free lock at 2^32^ and 2^31^, precision level L1 unless marked L2. n.a., not applicable; n.m., not measured.

| Model | Lock | Attacker | Attack | Training seeds | Held-out loss | Downstream AUROC |  |  |
|---|---|---|---|---|---|---|---|---|
|  |  |  |  |  | (bits/base) | Host Tropism | Pathogenicity | Transmissibility |
| Evo-2 | — | — | Pretrained (no fine-tuning) | — | 1.834 | 0.933 | 0.878 | 0.813 |
| Evo-2 | — | — | Unlocked fine-tuning | 9 | 1.566 | 0.939 | 0.899 | 0.842 |
| Evo-2 | SpecDef | Naive | Full fine-tuning | 3 | 1.893 | 0.902 | 0.874 | 0.644 |
| Evo-2 | SpecDef | Naive | Full fine-tuning, L2 | 3 | 1.942 | 0.711 | 0.799 | 0.601 |
| Evo-2 | SpecDef | Naive | LoRA on $\widetilde{W}$ | — | n.a. | 0.938 | 0.880 | 0.820 |
| Evo-2 | SpecDef | Naive | LoRA on $\widetilde{W}$ and *C* | — | n.a. | 0.565 | 0.583 | 0.568 |
| Evo-2 | SpecDef | Informed | Freeze *C* | 6 | 1.590 | 0.940 | 0.900 | 0.842 |
| Evo-2 | SpecDef | Informed | Freeze *C*, L2 | 3 | 1.564 | 0.934 | 0.901 | 0.847 |
| Evo-2 | SpecDef | Informed | Fuse $C\widetilde{W}$, before fine-tuning | — | 1.834 | 0.931 | 0.873 | 0.824 |
| Evo-2 | SpecDef | Informed | Freeze all 32 locked matrices | 3 | 1.574 | 0.938 | 0.896 | 0.826 |
| Evo-2 | Compensation-free | Naive | Full fine-tuning | 3 | 1.898 | 0.866 | 0.842 | 0.656 |
| Evo-2 | Compensation-free | Informed | Canonicalise, then fine-tune | 3 | 1.567 | 0.933 | 0.888 | 0.844 |
| Evo-1 | — | — | Pretrained (no fine-tuning) | — | 1.885 | 0.930 | 0.884 | 0.809 |
| Evo-1 | — | — | Unlocked fine-tuning | 3 | 1.665 | 0.940 | 0.892 | 0.842 |
| Evo-1 | — | — | Unlocked fine-tuning, fp32 AdamW | 3 | 1.669 | 0.945 | 0.897 | 0.837 |
| Evo-1 | SpecDef | Naive | Full fine-tuning | 3 | 1.986 | 0.672 | 0.752 | 0.575 |
| Evo-1 | SpecDef | Naive | Full fine-tuning, L2 | 3 | 1.990 | n.m. | n.m. | n.m. |
| Evo-1 | SpecDef | Informed | Freeze *C* | 3 | 1.678 | 0.944 | 0.896 | 0.842 |
| Evo-1 | SpecDef | Informed | Fuse $C\widetilde{W}$, before fine-tuning | — | 1.885 | n.m. | n.m. | n.m. |
| Evo-1 | Compensation-free | Naive | Full fine-tuning | 3 | 1.974 | 0.896 | 0.855 | 0.640 |
| Evo-1 | Compensation-free | Informed | Canonicalise, then fine-tune | 3 | 1.667 | 0.944 | 0.898 | 0.843 |

**Supplementary Table 4:** Naive attack across training settings (Evo-2). Held-out loss (bits per base) of the best trained checkpoint for unlocked fine-tuning and the naive attack on each lock, with SpecDef at *α* = 10^5^ and the compensation-free lock at 2^32^, precision level L1 and 6,000 steps unless stated. One training seed per cell, and three in the Adafactor and SGD rows (means). Every locked cell is significantly worse than pretrained, and every unlocked cell significantly better (genome bootstrap, 95%).

| Optimiser and learning rate | Unlocked | SpecDef | Compensation-free |
|---|---|---|---|
| AdamW (fp32 states), 10<sup>−7</sup> | 1.799 | 1.838 | 1.920 |
| AdamW (fp32 states), 3 × 10<sup>−7</sup> | 1.745 | 1.896 | 1.916 |
| AdamW (fp32 states), 10<sup>−6</sup> | 1.663 | 1.895 | 1.908 |
| AdamW (fp32 states), 3 × 10<sup>−6</sup> | 1.614 | 1.968 | 1.909 |
| AdamW (fp32 states), 10<sup>−5</sup> | 1.571 | 1.891 | 1.897 |
| AdamW (fp32 states), 3 × 10<sup>−5</sup> | 1.708 | 1.999 | 1.892 |
| AdamW (fp32 states), 10<sup>−4</sup> | 1.767 | 1.897 | 1.892 |
| AdamW (8-bit states), 10<sup>−5</sup>, no clipping | 1.654 | 1.983 | 1.955 |
| 24,000 steps, 10<sup>−5</sup> | 1.548 | 1.887 | 1.890 |
| 24,000 steps, 3 × 10<sup>−5</sup> | 1.698 | 1.885 | 1.888 |
| 24,000 steps, 10<sup>−4</sup> | 1.731 | 1.892 | 1.889 |
| Adafactor, 10<sup>−5</sup> | 1.579 | 1.892 | 1.901 |
| SGD with momentum 0.9, 10<sup>−3</sup> | 1.807 | 1.886 | 1.938 |

**Supplementary Table 5:**
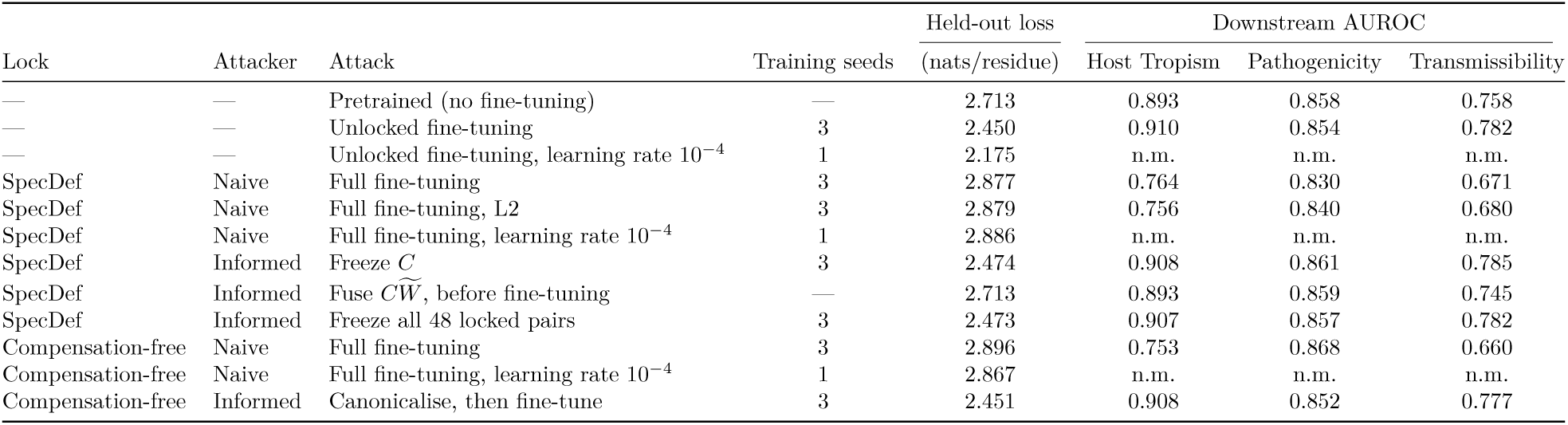
Held-out loss and downstream AUROC for each attack on ESM3-open. Held-out loss in nats per masked residue, weighted equally across held-out genomes, and AUROC at the best trained checkpoint. Values are means over training seeds, each averaged over three classifier seeds (six for the pretrained model). SpecDef at *α* = 10^5^ and compensation-free lock at 2^32^, precision level L1 and learning rate 10*^−^*^5^ unless stated. On Pathogenicity the naive attacks fall below pretrained only on the full test set. n.m., not measured.

|  |  |  |  | Held-out loss | Downstream AUROC |  |  |
|---|---|---|---|---|---|---|---|
| Lock | Attacker | Attack | Training seeds | (nats/residue) | Host Tropism | Pathogenicity | Transmissibility |
| — | — | Pretrained (no fine-tuning) | — | 2.713 | 0.893 | 0.858 | 0.758 |
| — | — | Unlocked fine-tuning | 3 | 2.450 | 0.910 | 0.854 | 0.782 |
| — | — | Unlocked fine-tuning, learning rate 10<sup>−4</sup> | 1 | 2.175 | n.m. | n.m. | n.m. |
| SpecDef | Naive | Full fine-tuning | 3 | 2.877 | 0.764 | 0.830 | 0.671 |
| SpecDef | Naive | Full fine-tuning, L2 | 3 | 2.879 | 0.756 | 0.840 | 0.680 |
| SpecDef | Naive | Full fine-tuning, learning rate 10<sup>−4</sup> | 1 | 2.886 | n.m. | n.m. | n.m. |
| SpecDef | Informed | Freeze *C* | 3 | 2.474 | 0.908 | 0.861 | 0.785 |
| SpecDef | Informed | Fuse $C\widetilde{W}$, before fine-tuning | — | 2.713 | 0.893 | 0.859 | 0.745 |
| SpecDef | Informed | Freeze all 48 locked pairs | 3 | 2.473 | 0.907 | 0.857 | 0.782 |
| Compensation-free | Naive | Full fine-tuning | 3 | 2.896 | 0.753 | 0.868 | 0.660 |
| Compensation-free | Naive | Full fine-tuning, learning rate 10<sup>−4</sup> | 1 | 2.867 | n.m. | n.m. | n.m. |
| Compensation-free | Informed | Canonicalise, then fine-tune | 3 | 2.451 | 0.908 | 0.852 | 0.777 |

